# IFNγ modestly contributes to disease progression in the *Ndufs4*(-/-) model of Leigh syndrome while IP10 is dispensable

**DOI:** 10.1101/2023.07.09.548281

**Authors:** Allison R Hanaford, Asheema Khanna, Katerina James, Vivian Truong, Ryan Liao, Yihan Chen, Michael Mulholland, Bernhard Kayser, Erin Shien Hsieh, Margaret Sedensky, Phil Morgan, Vandana Kalia, Surojit Sarkar, Simon C Johnson

## Abstract

Leigh syndrome (LS) is the most common pediatric presentation of genetic mitochondrial disease. LS is a multi-system disease characterized by severe neurologic and metabolic abnormalities. The defining feature of the disease is the presence of symmetric, bilateral, progressive necrotizing lesions in the brain stem, cerebellum, and basal ganglia. The pathogenic mechanisms underlying disease initiation and progression in LS have yet to be elucidated. Recent evidence demonstrates that the immune system plays a key role in LS pathogenesis. Treatment with the macrophage-depleting Csf1r inhibitor pexidartinib prevents disease in the *Ndufs4*(-/-) mouse model of LS, but the mechanisms leading to immune activation and governing disease progression remain to be elucidated. In recent work, the cytokines IFNγ and IFNγ-induced protein 10 (IP10) were found to be significantly elevated in *Ndufs4*(-/-) brainstem. Given their role as macrophage-activating factors, here we sought to assess the role of IFNγ and IP10 in LS using by generating *Ndufs4*(-/-)/*Ifng*(-/-) and *Ndufs4*(-/-)/*IP10*(-/-) double knockout lines. We find that IP10 alone does not significantly impact the onset or progression of disease in the *Ndufs4*(-/-) model, while IFNγ loss significantly, but modestly, improves survival. These data indicate that IFNγ contributes to pathology, but that IFNγ and IP10 are both dispensable for overall disease course of LS. Our findings support some role for IFNγ targeting therapies in the management of mitochondrial disease, but suggest they may provide only modest benefits, at least in LS.

## Introduction

Leigh syndrome (LS), also called subacute necrotizing encephalomyopathy, is the most common pediatric presentation of genetic mitochondrial disease [1, 2]. The primary pathological feature is symmetric, bilateral, necrotic lesions in the brainstem, basal ganglia and cerebellum [1, 3]. Patients are typically born healthy and develop symptoms of neurodegeneration in early childhood. Symptoms including dysphagia and respiratory issues result from brainstem dysfunction, which ultimately leads to death [1, 3]. LS is extremely genetically diverse, having been linked to more than 100 unique genetic defects appearing in both the nuclear and mitochondrial genomes [4].

Mutations in *NDUFS4* have been causally linked to LS in humans [5]. Homozygous deletion of *Ndufs4* in mice results in a phenotype remarkably consistent with human LS. As in human patients, *Ndufs4*(-/-) mice are born healthy and develop neurologic symptoms early in life, at around 5-6 weeks of age. These animals develop necrotic inflammatory lesions in the brainstem and cerebellum. In advanced disease, the lesions these enriched in IBA1(+) phagocytes. *Ndufs4*(-/-) mice have a significantly shortened lifespan, with a median survival of about 60 days [6-8].

Our lab recently demonstrated that disease in *Ndufs4*(-/-) mice is medicated by immune cells. Pharmacologic depletion of circulating and tissue macrophages (including brain microglia) with a high dose Csf1r (Colony stimulating factor 1 receptor) inhibitor prevents formation of necrotic lesions, prevents neurologic disease symptoms, and dramatically prolongs lifespan, which is limited by drug toxicity [9]. Csf1r is critical for the survival and function of macrophages, including brain resident microglia [10].

In this study, cytokine profiling revealed that interferon-gamma (IFNγ) and IFNγ-induced protein 10 (IP10, also known as Cxcl10) were significantly elevated in *Ndufs4*(-/-) mice with overt disease compared to age-matched control animals [9]. IFNγ has pleiotropic functions in the innate and adaptive immune systems. It is produced in response to viral infection and signals through the IFNγ receptor (IFNγR) which activates the JAK-STAT pathway leading to activation of interferon stimulated genes, including *IP10/CXCL10*. IP10 acts through the chemokine receptor CXCR3 and, among other functions, is known to act as a potent chemoattractant molecule for a variety of leukocytes including monocytes/macrophages, T cells, and NK cells [11, 12]. Given that neuroinflammatory lesions in LS are defined by the accumulation of mononuclear phagocytic cells, inhibition of Csf1r by pexidartinib prevents disease, and IFNγ and IP10 are significantly elevated in the brains of *Ndufs4*(-/-) mice, we sought to elucidate the roles of IFNγ and IP10 in LS. Here, we define the role of IFNγ and IP10 by genetically disrupting *Ifng* and *IP10* using knockout lines crossed into the *Ndufs4*(-/-) model.

## Results

### Loss of IP10 does not attenuate disease in the Ndufs4(-/-) mouse model of LS

To evaluate the role of IP10 in LS, we generated and assessed disease onset and progression in *Ndufs4*(-/-) animals with wildtype, heterozygous knockout, or homozygous knockout of *IP10* by crossing our *Ndufs4*(+/-) animals with the well-established *IP10* knockout line (Jackson laboratory strain 002287, see **Methods**).

Neither heterozygous nor homozygous loss of *IP10* had any significant impact on measures of health or neurologic disease in the *Ndufs4*(-/-) model (**Figure 1A-F**). IP10 loss did not impact body weight or the onset of weight loss (**Figure 1A-B**). Loss of *IP10* also failed to delay the onset of ataxia or forelimb clasping in the *Ndufs4*(-/-) model (**Figure 1C-E**). Consistent with these findings, the progressive decline in rotarod performance which occurs in the *Ndufs4*(-/-) model was not attenuated by loss of IP10 (**Figure 1F**).

**Figure 1.**
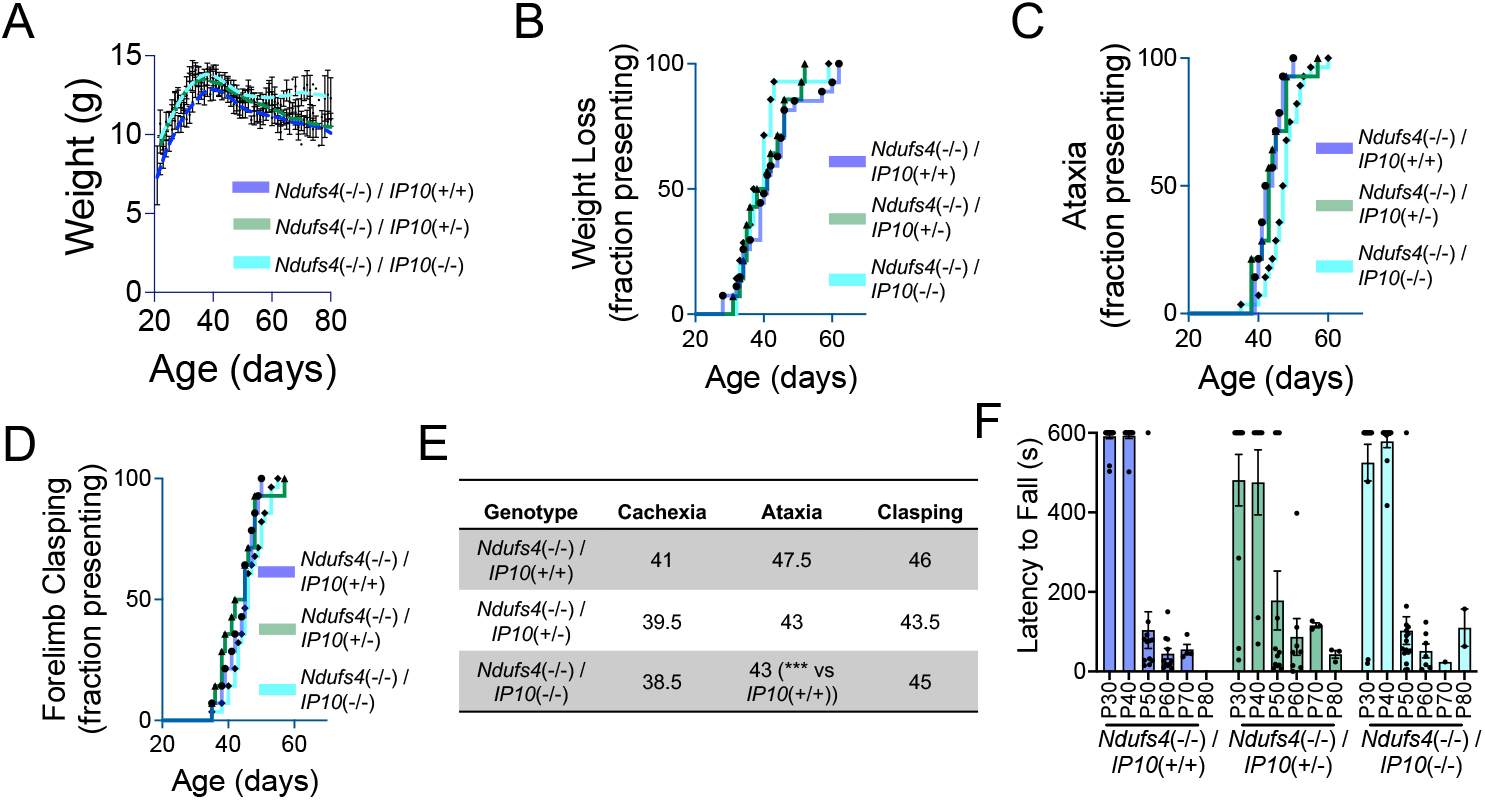
Loss of *IP10* does not impact disease course in *Ndufs4*(KO) mice. (A) Weight of *Ndufs4*(-/-)/*IP10*(+/+), *Ndufs4*(-/-)/*IP10*(+/-) and *Ndufs4*(-/-)/*IP10*(-/-) animals. Data shown are population averages with standard error of the mean (SEM) and Locally Weighted Scatterplot Smoothing (LOWESS) curves to show overall population trends. (B) Onset of weight loss (see *Methods*) in *Ndufs4*(-/-)/*IP10*(+/+), *Ndufs4*(-/-)/*IP10*(+/-) and *Ndufs4*(-/-)/*IP10*(-/-) animals. Pairwise log-rank test comparisons = not significant. (C) Onset of ataxia in *Ndufs4*(-/-)/*IP10*(+/+), *Ndufs4*(-/-)/*IP10*(+/-) and *Ndufs4*(-/-)/*IP10*(-/-) animals. Log-rank test ***p≤0.001 for *Ndufs4*(-/-)/*IP10*(+/+) *vs Ndufs4*(-/-)/*IP10*(-/-), other pairwise comparisons not significant. (D) Onset of forelimb clasping in *Ndufs4*(-/-)/*IP10*(+/+), *Ndufs4*(-/-)/*IP10*(+/-) and *Ndufs4*(-/-)/*IP10*(-/-) animals. Pairwise log-rank test comparisons = not significant. (E) Summary of median ages of symptom appearance. ***p<0.001 by Log-rank test, see (C). (F) Rotarod performance as assessed by latency to fall (see **Methods**). No significant comparisons by two-way ANOVA with Tukey multiple testing correction (all possible comparisons made) or by pairwise comparisons between genotypes at the same age.

**Figure 2.**
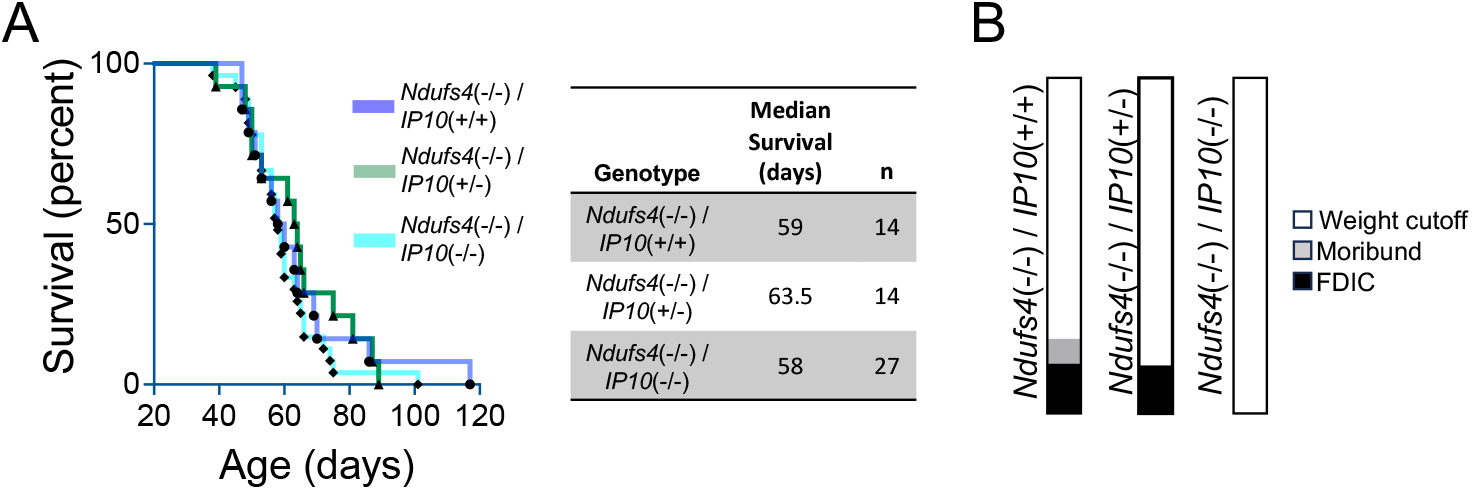
Loss of *IP10* does not improve survival in *Ndufs4*(KO) mice. (A) *Ndufs4*(-/-)/*IP10*(+/+), *Ndufs4*(-/-)/*IP10*(+/-) and *Ndufs4*(-/-)/*IP10*(-/-) animals. Summary of median survival and number of animals for each genotype cohort provided. No pairwise log-rank comparisons were statistically significant. (B) Cause of death in *Ndufs4*(-/-)/*IP10*(+/+), *Ndufs4*(-/-)/*IP10*(+/-) and *Ndufs4*(-/-)/*IP10*(-/-) animals.

Survival was not altered by IP10 loss in the *Ndufs4*(-/-) (**Figure 4A-B**), and the primary cause of death in *Ndufs4*(-/-) mice was euthanasia due to weight loss regardless of *IP10* status (**Figure 4C**). Together, the data indicate that *IP10* is dispensable in *Ndufs4*(-/-) disease onset and progression.

### Loss of IFNγ modestly impacts disease progression and survival in the Ndufs4(-/-) model

To evaluate the role of IFNγ in LS, we generated and assessed disease onset and progression in *Ndufs4*(-/-) animals with wildtype, heterozygous knockout, or homozygous knockout of *Ifng* by crossing our *Ndufs4*(+/-) animals with the well-established *Ifng* knockout line (Jackson laboratory strain 006087, see **Methods**).

Homozygous loss of *Ifng* resulted in a modest reduction in body size and delay in the onset of weight loss (**Figure 3A-B**). Clasping onset was also very modestly delayed by homozygous loss of *Ifng*, but the onset of ataxia was not affected (**Figure 3C-E**, see **Methods**). As in the Neurological function was assessed by observing onset of ataxia and forelimb clasping and the rotarod performance test. Homozygous or heterozygous loss of *Ifng* did not significantly alter the age of ataxia onset (**Figure 1C**) or forelimb clasping (**Figure 1D**). Rotarod performance appeared marginally improved by *Ifng* knockout at P50, but benefits were not maintained to later ages of P60 and 70 (**Figure 3F**).

**Figure 3.**
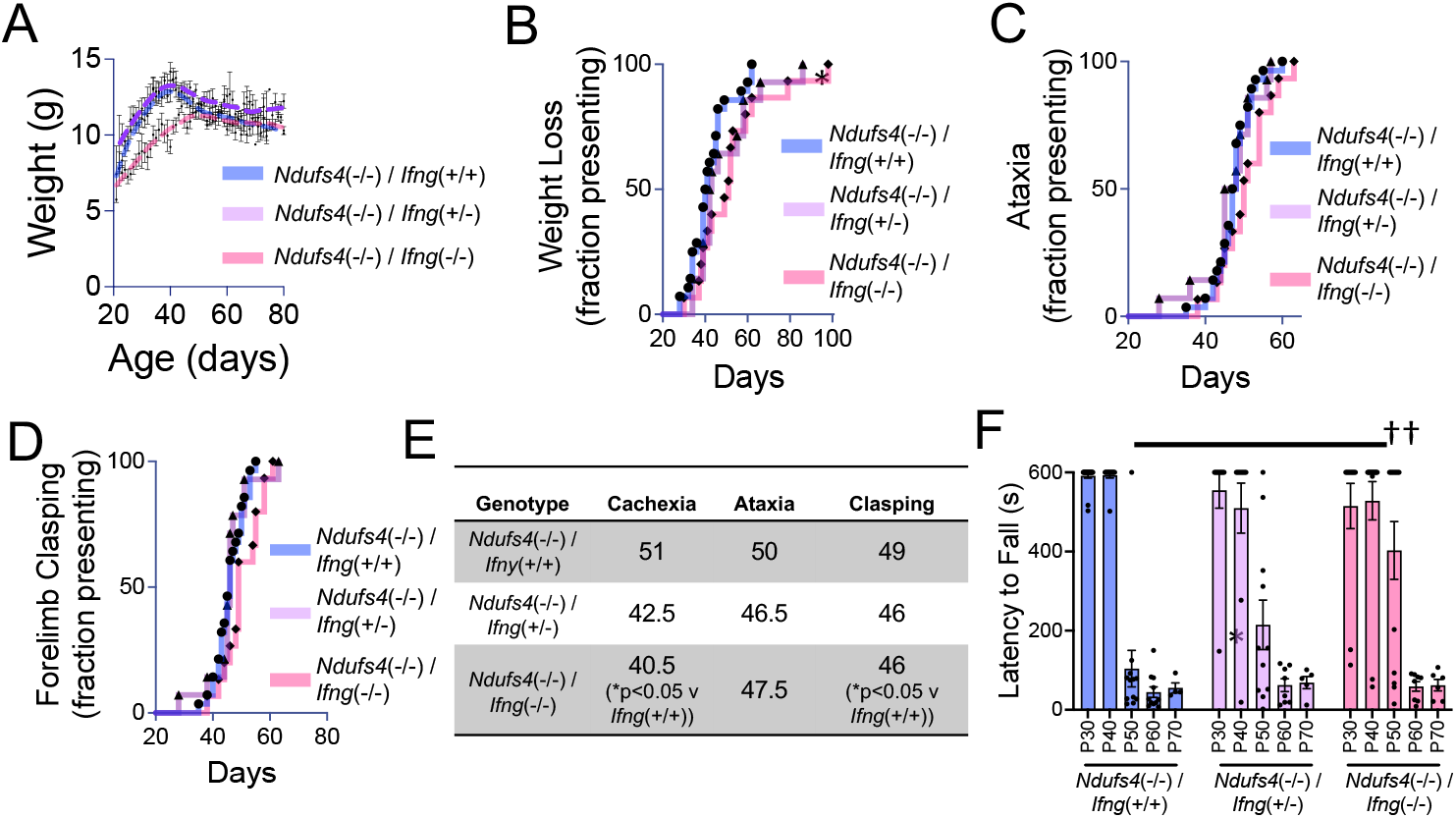
Loss of *Ifng* delays onset of weight loss and forelimb clasping in *Ndufs4*(KO) mice. (A) Average weight of *Ndufs4*(-/-)/*Ifng*(+/+), *Ndufs4*(-/-)/*Ifng*(+/-), and *Ndufs4*(-/-)/*Ifng*(-/-) animals. Data shown are population averages with standard error of the mean (SEM) and Locally Weighted Scatterplot Smoothing (LOWESS) curves to show overall population trends. (B) Onset of weight loss (see *Methods*) in *Ndufs4*(-/-)/*Ifng*(+/+), *Ndufs4*(-/-)/*Ifng*(+/-), and *Ndufs4*(-/-)/*Ifng*(-/-) animals. Log-rank test *p<0.05 for *Ndufs4*(-/-)/*IP10*(+/+) *vs Ndufs4*(-/-)/*IP10*(-/-), other pairwise comparisons not significant. (C) Onset of ataxia in *Ndufs4*(-/-)/*Ifng*(+/+), *Ndufs4*(-/-)/*Ifng*(+/-), and *Ndufs4*(-/-)/*Ifng*(-/-) animals. Log-rank comparisons not significant. (D) Onset of forelimb clasping in *Ndufs4*(-/-)/*Ifng*(+/+), *Ndufs4*(-/-)/*Ifng*(+/-), and *Ndufs4*(-/-)/*Ifng*(-/-) animals. Log-rank test *p<0.05 for *Ndufs4*(-/-)/*IP10*(+/+) *vs Ndufs4*(-/-)/*IP10*(-/-), other pairwise comparisons not significant. (E) Summary of median ages of symptom appearance. *p<0.05 by Log-rank test. (F) Rotarod performance as assessed by latency to fall (see **Methods**). No significant comparisons by two-way ANOVA with Tukey multiple testing correction (all possible comparisons made). ††p<0.005 by pairwise T-test between *Ndufs4*(-/-)/*IP10*(+/+) and *Ndufs4*(-/-)/*IP10*(-/-) at P50.

While the overall impact of *Ifng* loss on the onset of neurologic symptoms was marginal, *Ifng* loss resulted in a gene dosage-dependent increase in survival: median survival was 70.5 and 86 days, respectively, in *Ndufs4*(-/-)/*Ifng*(+/-) and *Ndufs4*(-/-)/*Ifng*(-/-) compared to 58 days in the *Ndufs4*(-/-)/*Ifng*(+/+) cohort (**Figure 4A-B)**. Survival was significantly increased by both heterozygous and homozygous loss of *Ifng*. The primary cause of death in *Ndufs4*(-/-) mouse, regardless of *Ifng* status, was euthanasia due to weight loss (**Figure 4C**).

**Figure 4.**
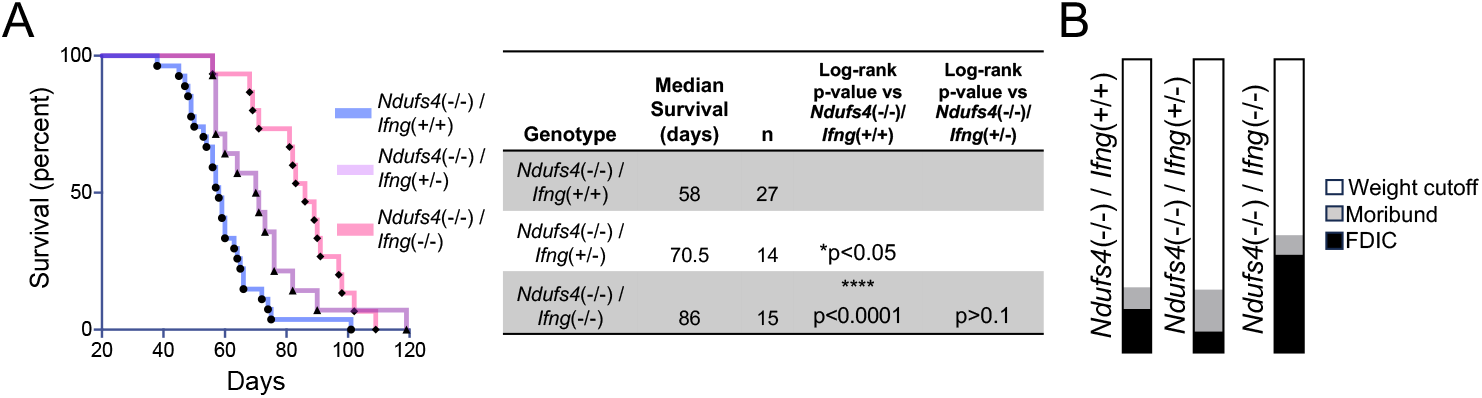
Loss of *Ifng* extends survival in *Ndufs4*(KO) mice. (A) Survival of *Ndufs4*(-/-)/*Ifng*(+/+), *Ndufs4*(-/-)/*Ifng*(+/-), and *Ndufs4*(-/-)/*Ifng*(-/-) animals. Summary of median survival and cohort numbers by genotype and p<values of pairwise group comparisons by Log-rank test. (B) Cause of death in *Ndufs4*(-/-)/*Ifng*(+/+), *Ndufs4*(-/-)/*Ifng*(+/-), and *Ndufs4*(-/-)/*Ifng*(-/-) animals.

## Discussion

Here, we used genetic models to assess the role of IP10 and IFNγ, two inflammatory cytokines previously found to be significantly elevated in diseased *Ndufs4*(-/-) brainstem tissue, in the onset and progression of disease in the *Ndufs4*(-/-) model of LS. Our data indicate that the leukocyte chemoattractant IP10 is dispensable for both disease onset and progression, while the IFNγ, an upstream regulator of IP10 and important factor in both innate and adaptive immunity, contributes modestly but significantly to weight loss, survival, and the onset of clasping and impaired rotarod performance.

While the immunologic origins of disease in LS were demonstrated in our prior study, the specific immunological factors driving leukocyte driven brainstem lesion formation, as well as the upstream immune-activating signals, remain undefined. We chose to define the roles of IP10 and IFNγ based on cytokine profiling from the *Ndufs4*(-/-), where they were found to be significantly elevated [9], and a number of recent studies linking IFNγ to mitochondrial dysfunction. In particular, IFNγ has been shown to be significantly elevated in patients with multiple genetically distinct forms of mitochondrial dysfunction, as well as in age-related mitochondrial dysfunction, and is thought to mediate mitochondrial dysfunction associated immune-activation in other settings [4, 11, 13-17]. Our data here indicate that IFNγ does indeed participate in disease, but neither IFNγ, nor the IFNγ factor IP10, are necessary for the overall pathobiology of LS.

Our findings both provide additional support for the immune-mediated model of LS pathogenesis and demonstrate that IFNγ contributes only partially to disease progression in the *Ndufs4*(-/-) model. These findings move us one step closer to an understanding of the immune mediators responsible for pathology in settings of genetic mitochondrial disease.

## Methods

### Animals

*IP10*(-/-) and *Ifng*(-/-) mice are from the Jackson Lab (strains 002287 and 006087). *Ndufs4*(+/-) mice were originally obtained from the Palmiter laboratory at University of Washington, Seattle, Washington USA, but are also available from the Jackson Laboratory (strain 027058). Strain details are described in Kruse et al [7]. *Ndufs4*(-/-), *Ifng*(-/-), and *IP10*(-/-) lines are all on the C57/BL6 background. *Ndufs4*(-/-) mice cannot be used for breeding due to their short lifespan and severe disease. *Ndufs4*(+/-) mice were bred with *Ifng*(-/-) or *IP10*(-/-) mice to produce double heterozygous offspring, which were then crossed to produce *Ndufs4*(-/-)/*Ifng*(+/-), *Ndufs4*(-/-)/*Ifng*(-/-), *Ndfus4*(-/-)/*IP10(+/-*), and *Ndufs4*(-/-)/*IP10*(-/-) offspring. Genotyping of the *Ndufs4, Ifng*, and *IP10* alleles were performed according to the Jackson laboratory methods (strains 027058, 002287, and 006087 respectively)

Mice were weaned at P20-22 days of age. *Ndufs4*(-/-) animals were housed with control littermates for warmth as *Ndufs4*(-/-) mice have low body temperature [7]. Mice were weighed and health assessed a minimum of 3 times a week. Following onset of *Ndufs4*(-/-) symptoms, wet food was provided in the bottom of the cage. Animals were euthanized if they reached a 20% loss of maximum body weight (measured two days consecutively), were immobile, or found moribund. Mice heterozygous for *Ndufs4* have no reported phenotype, so controls consisted of both heterozygous and wild-type *Ndufs4* animals. The *Ndufs4*(Ctrl) and *Ndufs4*(-/-) mice wild type for *Ifng* or *IP10* used in this study came from crosses of *Ndufs4*(+/-)/*Ifng*(+/-) mice, *Ndufs4*(+/-)/*Ip10*(+/-) mice, and our general Ndufs4 colony. Mice were fed PicoLab Diet 5058 and were on a 12-hour light-dark cycle. All animal experiments followed Seattle Children’s Research Institute (SCRI) guidelines and were approved by the SCRI IACUC.

Clasping, circling, and ataxia were assessed by visual scoring and analyzed as previously described [8]. During disease progression *Ndufs4*(-/-) animals can display intermittent/transient improvement of symptoms, so here we report whether the animal *ever* displayed the symptoms for two or more consecutive days.

The rotarod performance test was performed using a Med Associates ENV-571M single-lane rotarod. A mouse was placed on the rod already rotating at 6 rpm and latency to fall was timed for a maximum of 600 seconds while rotation remained constant. For each mouse, three trials were performed with a minimum of 10 minutes between each trial. The best of three trials was reported.

### Statistical analysis

All statistical analyses were performed using GraphPad prism, with statistical tests detailed in figure legends. Error bars represent the standard error of the mean (SEM).

### Scientific rigor

*Sex-*Both male and female animals were used in these experiments. No significant sex differences have been reported in *Ndfus4(-/-), Ifng*(-/-), or *IP10*(-/-) and none were observed.

*Exclusion criteria-*Animals euthanized prior to the age of disease onset in the *Ndufs4*(-/-) were excluded from study. Our criteria for early life exclusion includes severe weaning stress (significant weight loss or spontaneous mortality before P30), runts (defined as ≤5 g body weight at weaning age), or those or born with health issues unrelated to the *Ndufs4*(-/-) phenotype (such as hydrocephalus). These criteria are applied to all genotypes as part of our standard animal care.

